# Genomic surveillance for hypervirulence and multi-drug resistance in invasive *Klebsiella pneumoniae* from south and southeast Asia

**DOI:** 10.1101/557785

**Authors:** Kelly L Wyres, To N T Nguyen, Margaret M C Lam, Louise M Judd, Nguyen van Vinh Chau, David A B Dance, Margaret Ip, Abhilasha Karkey, Clare L Ling, Thyl Miliya, Paul N Newton, Lan Nguyen, Amphone Sengduangphachanh, Paul Turner, Balaji Veeraraghavan, Phat Voong Vinh, Manivanh Vongsouvath, Nicholas R Thomson, Stephen Baker, Kathryn E Holt

## Abstract

**Background:** *K. pneumoniae* is a leading cause of blood stream infection (BSI). Strains producing extended spectrum beta-lactamases (ESBLs) or carbapenemases are considered global priority pathogens for which new treatment and prevention strategies are urgently required, due to severely limited therapeutic options. South and Southeast Asia are major hubs for antimicrobial resistant (AMR) *K. pneumoniae*, and also for the characteristically antimicrobial sensitive, community-acquired ‘hypervirulent’ strains. The emergence of hypervirulent AMR strains and lack of data on exopolysaccharide diversity pose a challenge for *K. pneumoniae* BSI control strategies worldwide.

**Methods:** We conducted a retrospective genomic epidemiology study of 365 BSI *K. pneumoniae* from seven major healthcare facilities across South and Southeast Asia, extracting clinically relevant information (AMR, virulence, K and O antigen loci) using *Kleborate*.

**Findings:** *K. pneumoniae* BSI isolates were highly diverse, comprising 120 multi-locus sequence types (STs) and 63 K-loci. ESBL and carbapenemase gene frequencies were 47% and 17%, respectively. The aerobactin synthesis locus (*iuc*), associated with hypervirulence, was detected in 28% of isolates. Importantly, 7% of isolates harboured *iuc* plus ESBL and/or carbapenemase genes. The latter represent genotypic AMR-virulence convergence, which is generally considered a rare phenomenon but was particularly common amongst South Asian BSI (17%). Of greatest concern, we identified seven novel plasmids carrying both *iuc* and AMR genes, raising the prospect of co-transfer of these phenotypes amongst *K. pneumoniae*.

**Interpretation:** South and Southeast Asia are high-risk regions for the emergence of AMR and convergent AMR-hypervirulent *K. pneumoniae*. Enhanced surveillance efforts, reporting STs, AMR and virulence information are urgently required to monitor this public health threat.

**Funding:** This work was supported by the Wellcome Trust (grant #206194 to Wellcome Sanger Institute) and the Bill and Melinda Gates Foundation, Seattle (grant OPP1175797 to KEH). KEH is supported by a Senior Medical Research Fellowship from the Viertel Foundation of Australia. DAB and PNN are supported by the Wellcome Trust.

## Background

*Klebsiella pneumoniae* is now regarded globally by the World Health Organisation (WHO) and others, as a priority antimicrobial resistant (AMR) pathogen requiring new control strategies^1^. These include rapid identification and containment of high-risk AMR clones such as the carbapenemase-producing (CP) variants, augmented with vaccines, bacteriophages, or immunotherapies that target conserved surface antigens. However, *K*. *pneumoniae* is highly diverse, hindering the development of such strategies and our ability to study its molecular epidemiology in a short time frame.

This diverse bacterial species is generally associated with a range of differing community and healthcare-associated infections, but can be particularly problematic when the organisms gain access to sterile sites such as the cerebrospinal fluid, internal body cavities, and the bloodstream. Such infections are often characterised by rapid onset and multi-drug resistance (MDR), including resistance to third generation cephalosporins and/or carbapenems. Antimicrobials are the primary treatment strategy but options are severely limited by AMR. Concomitant with this are elevated mortality rates and treatment costs^2^.

*K. pneumoniae* is amongst the most common cause of bloodstream infections (BSI) in South (S) and Southeast (SE) Asia^3–5^. Infections in these locations are associated with a high mortality rate^3, 6^, yet *K. pneumoniae* is generally understudied in Asia and we have limited insight into the population structure and diversity of the organisms causing these infections in this global AMR epicentre. Available data suggest a heterogenous landscape; for example, CP strains are rare in SE Asia (<1-4%^4, 7, 8^) but common in S Asia (28-70%^9, 10^) and the prevalence of extended-spectrum beta-lactamase (ESBL) producing organisms varies from 12-79% in these regions^3–5, 7, 9^. Studies investigating ESBL and CP variants in S/SE Asia implicate CTX-M-15 as the most common ESBL type^11^, while NDM and OXA-48-like enzymes are the most commonly described carbapenemases^10, 12^.

Whilst the picture of *K. pneumoniae* pathogenesis is incomplete, isolates harbouring one or more key virulence determinants (including RmpA/RmpA2, which upregulate capsule expression; the colibactin genotoxin; and the yersiniabactin, aerobactin and salmochelin siderophores that promote systemic survival and dissemination^13^) are more commonly associated with invasive disease^14, 15^. Isolates that contain several of these determinants are associated with severe community-acquired invasive disease, often manifesting as liver abscess with bacteraemia. These are classified as ‘hypervirulent’ infections, they occur globally but are most commonly reported in SE Asia^16^ and are associated with clones ST23, ST86, and ST65. These clones typically express the K1/K2 capsules and are generally susceptible to most antimicrobials. However, the last few years have seen increasing reports of ‘convergent’ *K. pneumoniae* that are both hypervirulent (carrying the *iuc* aerobactin locus, which is suggested to be the single most important feature of hypervirulent strains^17^) and MDR ESBL/carbapenemase producers. The majority of these reports represent sporadic isolations, but in 2017 Gu and colleagues reported an outbreak in a Chinese hospital that caused five deaths^18^. Given the high burden of both MDR and hypervirulent *K. pneumoniae* infections, S/SE Asia likely represents a major global hub for phenotypic convergence, with the potential for outbreaks of invasive disease with severely limited treatment options.

Here we present a genomic epidemiology study of BSI *K. pneumoniae* from seven major healthcare facilities across S/SE Asia, leveraging a recently established genomic framework that incorporates rapid genotyping of clinically important features (ST or ‘clone’, AMR, virulence, capsule and lipopolysaccharide serotypes). The data are highly relevant to the design of *K. pneumoniae* control strategies, revealing a diverse population with high rates of AMR and virulence loci, and high prevalence of potentially dangerous convergent strains.

## Methods

### Setting

The following tertiary care hospitals were included: Patan Hospital, Kathmandu, Nepal; Christian Medical College Hospital, Vellore, India; Angkor Hospital for Children, Siem Reap, Cambodia; Mahosot Hospital, Vientiane, Lao PDR; The Hospital of Tropical Diseases, Ho Chi Minh City, Vietnam; Prince of Wales Hospital, Hong Kong. Isolates were also provided from healthcare inpatient departments serviced by the Shoklo Malaria Research Unit, Mae Sot, Thailand.

### Bacterial isolates and culture

*K. pneumoniae* isolates obtained from blood cultures, following routine diagnostic protocols in each hospital laboratory and identified using biochemical testing (typically API-20E, bioMerieux), were included in the study. Isolates available for sequencing represented the following fractions of *K. pneumoniae* BSI collected at each site in participating years: India, 10%; Hong Kong, 17%; Vietnam, 20%; other sites, >90% (**Figure 1**).

**Figure 1:**
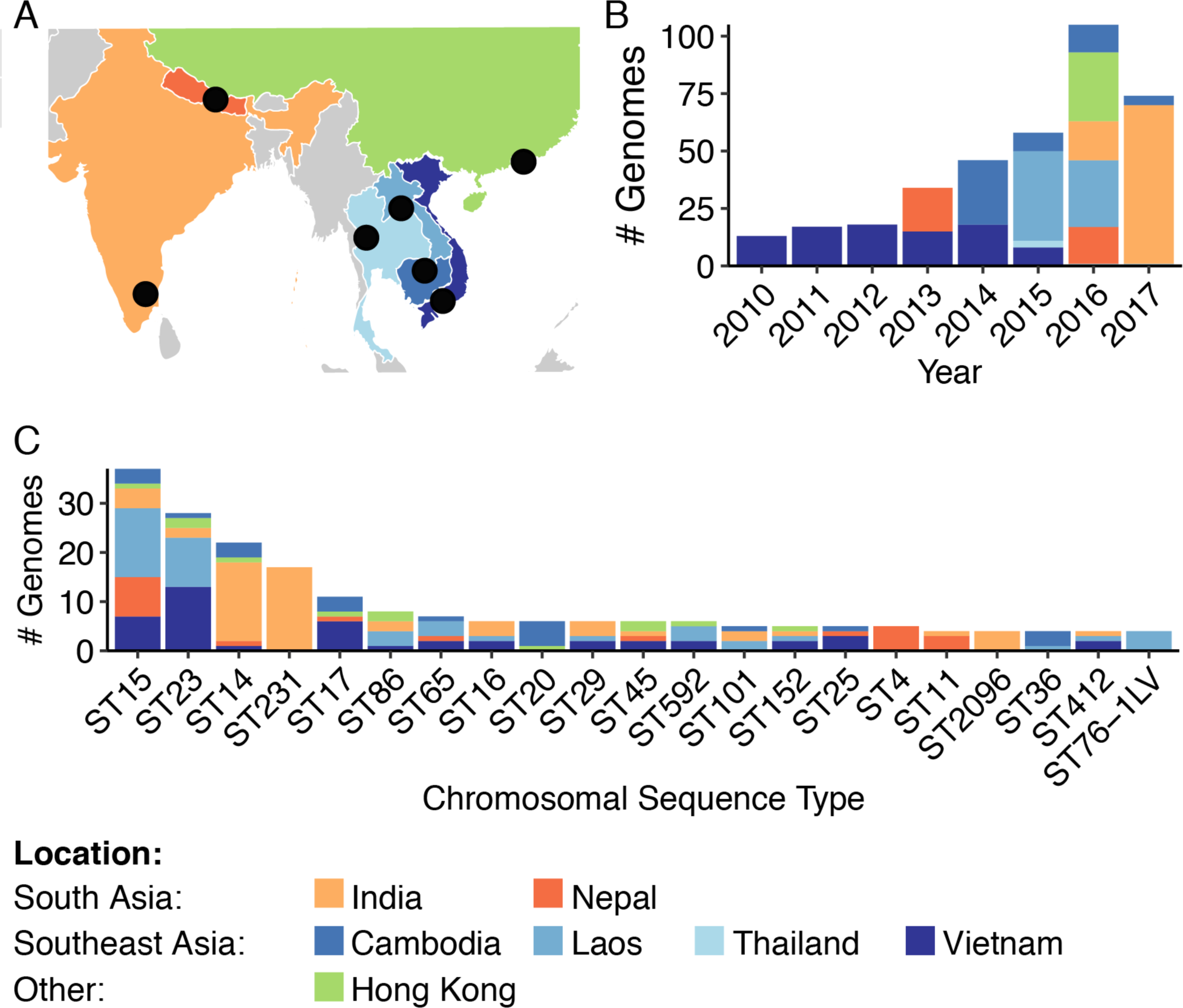
*Klebsiella pneumoniae* BSI isolates included in this study. **A)** Collection sites and countries of origin for all *Klebsiella pneumoniae* complex isolates for which genome data were available. **B)** Years of collection coloured by country of origin as in panel A. **C)** Chromosomal multi-locus sequence types (STs) of *Klebsiella pneumoniae sensu stricto* isolates (only STs accounting for >1% isolates are shown, coloured by country of origin as in panel A).

### DNA extractions, library preparation and sequencing

All isolates were cultured in LB broth at 37℃ overnight before DNA extraction using either Qiagen (Cambodia) or Promega kits (all other sites), as per manufacturer’s guidelines. Multiplexed Nextera XT libraries were sequenced on Illumina platforms, generating 150 bp paired-end reads. Eight isolates were selected for additional long-read sequencing using the Nanopore MinION R9 device as previously described^19^.

### Genome assembly and genotyping

Illumina adapter sequences were removed and reads were quality trimmed using TrimGalore v0.4.4 (https://github.com/FelixKrueger/TrimGalore). Subsequently, draft *de novo* assemblies were generated using SPAdes v3.10.1^20^ optimised with Unicycler v0.4.7^21^. We excluded from further analysis nine low-quality genome assemblies outside the expected size range (5–6.5 Mbp).

Chromosomal multi-locus sequence typing (MLST), virulence locus and acquired resistance genes (excluding the core ampicillin resistance gene *bla*_SHV_ and *oqxAB* efflux genes) were typed using *Kleborate* v0.3.0 (https://github.com/katholt/Kleborate). Surface capsule (K) and lipopolysaccharide (O) loci were identified using *Kaptive*^22, 23^. Novel K-loci were manually extracted using Bandage^24^, annotated using Prokka v1.13.3^25^ before manual curation, and deposit in GenBank (accessions TBD). Where *Kaptive* was unable to confidently identify a K- locus due to fragmented genome assemblies (n=123), K-loci were predicted from *wzi* alleles if an unambiguous locus was associated to the allele in the *K. pneumoniae* BIGSdb (https://bigsdb.pasteur.fr/klebsiella/klebsiella.html). Ska v1.0^26^ was used to calculate pairwise nucleotide differences between assemblies, and identify clusters of isolates for which the genomes differed by ≤100 single nucleotide variants (SNVs).

Hybrid Illumina-Nanopore assemblies were generated using Unicycler v0.4.7 as described previously^19^. Assemblies were annotated using Prokka v1.13.3 and *iuc+* plasmid annotations were manually curated before depositing in GenBank (accessions TBD). Plasmid replicon types were identified using the PlasmidFinder database v2.0^27^.

Isolate information, genotypes and genome data accessions are provided in **Table S1**. Genome assemblies, novel plasmid and K-locus sequences are also available in Figshare: https://doi.org/10.26180/5c67982956721.

### Statistical analyses

Statistical analyses were performed using R v3.3.3 and data were visualised using ggplot2 v2.2.1. Given the disparity in sampling frames and small subgroup sample sizes it was not appropriate to test trends at country or year level. Statistical tests for regional differences in genetic features between *K. pneumoniae* populations (**Table 1**) were calculated using the subset of *K. pneumoniae* collected in overlapping two-year periods: S Asia region (Nepal and India; 2016-2017; n=102) vs SE Asia region (Cambodia, Laos, Thailand, Vietnam; 2015-2016, n=100).

**Table 1.** Comparison of key features of Klebsiella genomes from South and South East Asian sites. South (S) Asia is represented by the two sites in India and Nepal, isolated 2016- 2017; South East (SE) Asia is represented by the sites in Cambodia, Laos, Thailand, Vietnam, isolated 2015-2016. P-values were calculated using Fishers Exact test and adjusted using Bonferroni correction for the number of tests within each group of comparisons (as labelled in bold; sample size for each are also in bold). Na; test not applicable (one or more values equal to zero). *p <0.05, **p <0.01

|  | S Asia | SE Asia | p-value (adj) | OR |  |
| --- | --- | --- | --- | --- | --- |
| **Species assignment** | **102** | **100** |  |  |  |
| *Klebsiella pneumoniae* | 100 (98%) | 87 (87%) | 0.011 | 0.13 | * |
| *Klebsiella variicola* | 1 (1%) | 4 (4%) | 0.838 | 4.18 |  |
| *Klebsiella quasipneumoniae* subsp. *quasipneumoniae* | 1 (1%) | 0 (0%) | na | na |  |
| *Klebsiella quasipneumoniae* subsp. *similipneumoniae* | 0 (0%) | 9 (9%) | na | na |  |
| **Features of *K. pneumoniae* sensu stricto** |  |  |  |  |  |
| **Sequence type assignment** | **100** | **87** |  |  |  |
| ST15 | 11 (11%) | 15 (17%) | 1.158 | 1.68 |  |
| ST14 | 16 (16%) | 2 (2%) | 0.008 | 0.12 | ** |
| ST23 | 2 (2%) | 12 (14%) | 0.015 | 7.76 | * |
| ST231 | 17 (17%) | 0 (0%) | 5.14 x 10<sup>-5</sup> | 0.00 | ** |
| **Capsular serotype prediction** | **79** | **87** |  |  |  |
| KL1 | 2 (2.5%) | 13 (15%) | 0.012 | 6.70 | * |
| KL2 | 4 (5.1%) | 11 (13%) | 0.216 | 2.70 |  |
| **Antimicrobial resistance prediction** | **100** | **87** |  |  |  |
| Multidrug resistant (≥3 acquired classes) | 81 (81%) | 44 (51%) | 1.06 x 10<sup>-4</sup> | 0.24 | ** |
| Aminoglycosides | 76 (76%) | 41 (47%) | 0.001 | 0.28 | ** |
| Fluoroquinolones | 80 (80%) | 39 (45%) | 7.13 x 10<sup>-6</sup> | 0.21 | ** |
| Phenicol | 27 (27%) | 25 (29%) | 7.833 | 1.09 |  |
| Sulfonamides | 63 (63%) | 42 (48%) | 0.493 | 0.55 |  |
| Tetracyclines | 24 (24%) | 39 (45%) | 0.029 | 2.56 | * |
| Trimethoprim | 62 (62%) | 38 (44%) | 0.118 | 0.48 |  |
| 3rd Generation Cephalosporins (ESBL) | 60 (60%) | 41 (47%) | 0.949 | 0.60 |  |
| Carbapenems | 47 (47%) | 1 (1%) | 7.13 x 10<sup>-13</sup> | 0.01 | ** |
| **Virulence prediction** | **100** | **87** |  |  |  |
| Yersiniabactin (*ybt*) | 65 (65%) | 46 (53%) | 0.205 | 0.61 |  |
| Aerobactin (*iuc*) | 27 (27%) | 32 (37%) | 0.319 | 1.57 |  |
| ***iuc* lineages** | **100** | **87** |  |  |  |
| *iuc* 1 | 13 (13%) | 27 (31%) | 0.012 | 2.99 | * |
| *iuc* 3 | 0 (0%) | 5 (6%) | 0.061 | na |  |
| *iuc* 5 | 12 (12%) | 0 (0%) | 0.001 | 0.00 | ** |
| **AMR and virulence convergence** | **100** | **87** |  |  |  |
| ESBL and *ybt* | 45 (45%) | 26 (30%) | 0.144 | 0.52 |  |
| ESBL and *iuc* | 16 (16%) | 1 (1%) | 0.001 | 0.06 | ** |
| CP and *ybt* | 37 (37%) | 1 (1%) | 1.79 x 10<sup>-10</sup> | 0.02 | ** |
| CP and *iuc* | 13 (13%) | 0 (0%) | na | na |  |

## Results

The genomes of 393 presumptive *K. pneumoniae* BSI from seven countries across S/SE Asia were successfully sequenced (**Figure 1A, Table S1**). Twenty-eight genomes were identified as Enterobacteriaceae species outside of the *K. pneumoniae* species complex and were excluded from further analyses. Among the remaining 365 isolates, the majority (n=331, 91%) of organisms were confirmed to be *K. pneumoniae*. Among the regional comparator samples, these accounted for a higher proportion of genomes from S than SE Asia (98% vs 87%, p=0.011; see **Table 1**). The remaining isolates (detailed in **Table S2**) were *K. quasipneumoniae* subsp*. similipneumoniae* (n=20, 5.5%), *K. variicola* (n=9, 2.5%), and *K. quasipneumoniae* subsp. *quasipneumoniae* (n=5, 1.4%), which are not distinguishable from *K. pneumoniae* by standard microbiology methods^14, 28^.

The 331 *K. pneumoniae* were highly diverse and comprised 120 individual STs (Simpson’s diversity index=0.97), the majority (61%) of which were represented by a single isolate. Nevertheless, we observed four common STs that each accounted for >5% of the sequenced organisms: ST15 (n=37, 11%), ST23 (n=28, 8.5%), ST14 (n=22, 6.6%) and ST231 (n=17, 5.1%; **Figure 1C**). Amongst these, only ST15 was common across all sites, whereas ST23 was significantly associated with SE Asia (p=0.015) and ST14 and ST231 were both significantly associated with S Asia (p<0.01; see **Table 1**).

### Predicted capsular (K) and O antigen serotypes

In this collection of invasive *K. pneumoniae* from BSI we detected 63 different K-loci including four novel loci designated KL162-165 (Simpson’s diversity index=0.95), the majority (67%) of these K-loci were found in ≤3 *K. pneumoniae* isolates each. The most common K-loci were KL1 (n=31; including 28 ST23), KL2 (n=27; numerous STs including ST14, ST25, ST65 and ST86), KL51 (n=23; including 17 ST231) and KL24 (n=20; including 17 ST15), together accounting for 28% of *K. pneumoniae* BSI (**Figure 2**). Saliently, while ST23 and ST231 were each associated with only a single K-locus (KL1 and KL51, respectively), the two other most common STs were each associated with multiple K-loci (ST15: KL24=17, KL112=9, KL10=3, KL62=2, KL19=2; ST14: KL64=7, KL2=6, KL157=1), as has been previously observed for ST258^23^. Ten of the 12 previously described O-antigen encoding loci were also detected (Simpson’s diversity index=0.80). Loci predicted to encode serotypes O1 and O2 were the most common, together accounting for 71% of *K. pneumoniae* BSI (**Figure 2**). Notably, the O3b locus, which is considered to be rare^29^, was detected here at 8% prevalence across sites (mean 7% per site, range 0-11%).

**Figure 2:**
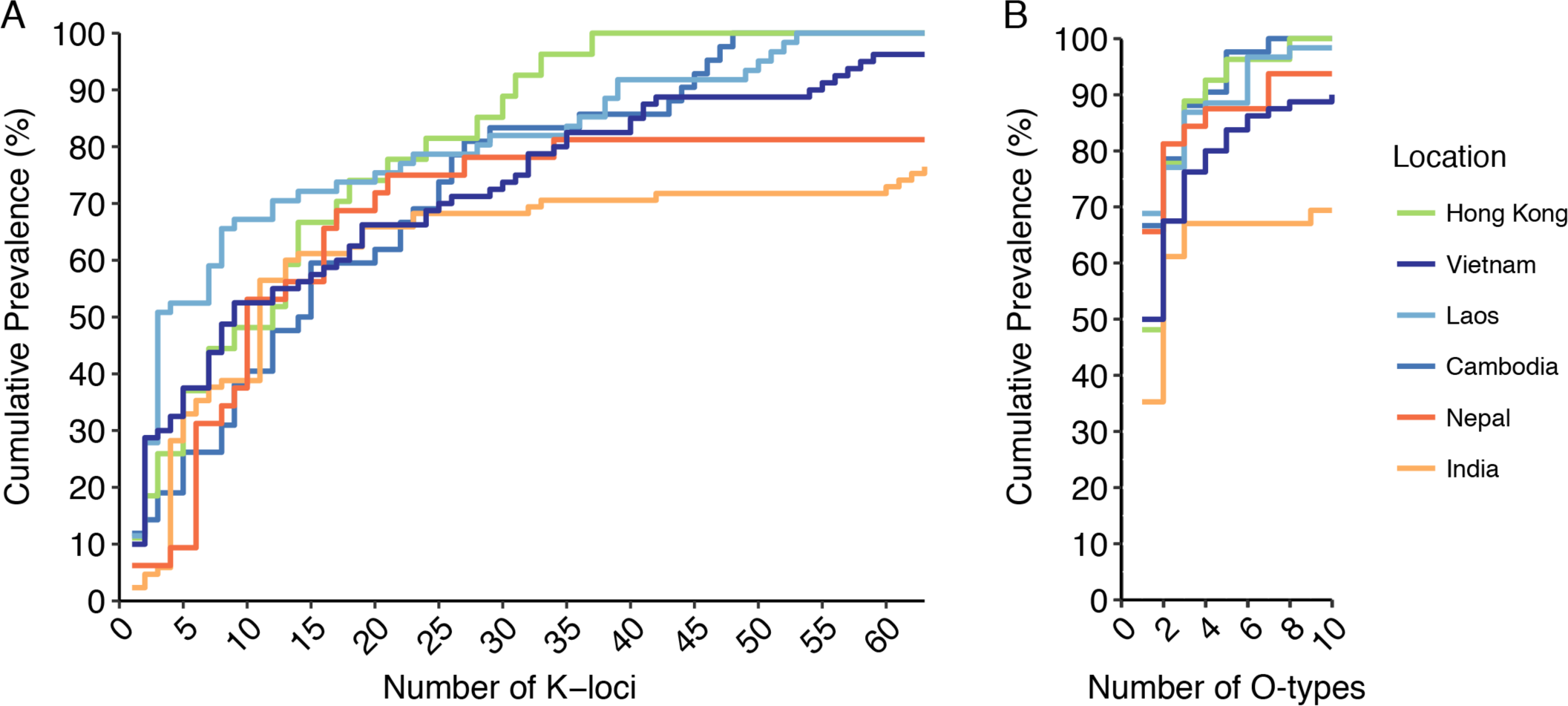
Cumulative K and O locus prevalence by location. **A)** Cumulative prevalence of K- loci ordered by mean prevalence across all locations (highest to lowest, see Table S3). **B)** Cumulative prevalence of predicted O-types ordered by mean prevalence across all locations (highest to lowest, see Table S4). Note that the Thai sampling site is excluded due to small sample size (n=4).

### Acquired antimicrobial resistance determinants

Acquired AMR determinants were detected in 91% of *K. pneumoniae* genomes. The number of antimicrobial classes to which each isolate was predicted to be resistant showed a bimodal distribution (**Figure 3A**), with the majority of *K. pneumoniae* being MDR (acquired AMR genes conferring resistance to ≥3 drug classes; 63%) or possessing no acquired AMR genes (9%). The prevalence of MDR differed between sampling location and ranged from 22% in Hong Kong to 85% in India. MDR was significantly more prevalent among S Asian isolates than SE Asian isolates (81% vs 51%, p=0.0001; see **Table 1**). Consistently, the S Asian organisms had a significantly higher prevalence of fluoroquinolone, aminoglycoside and carbapenem resistance determinants than the SE Asian organisms (*p*<0.001 for each class; see **Table 1**).

**Figure 3:**
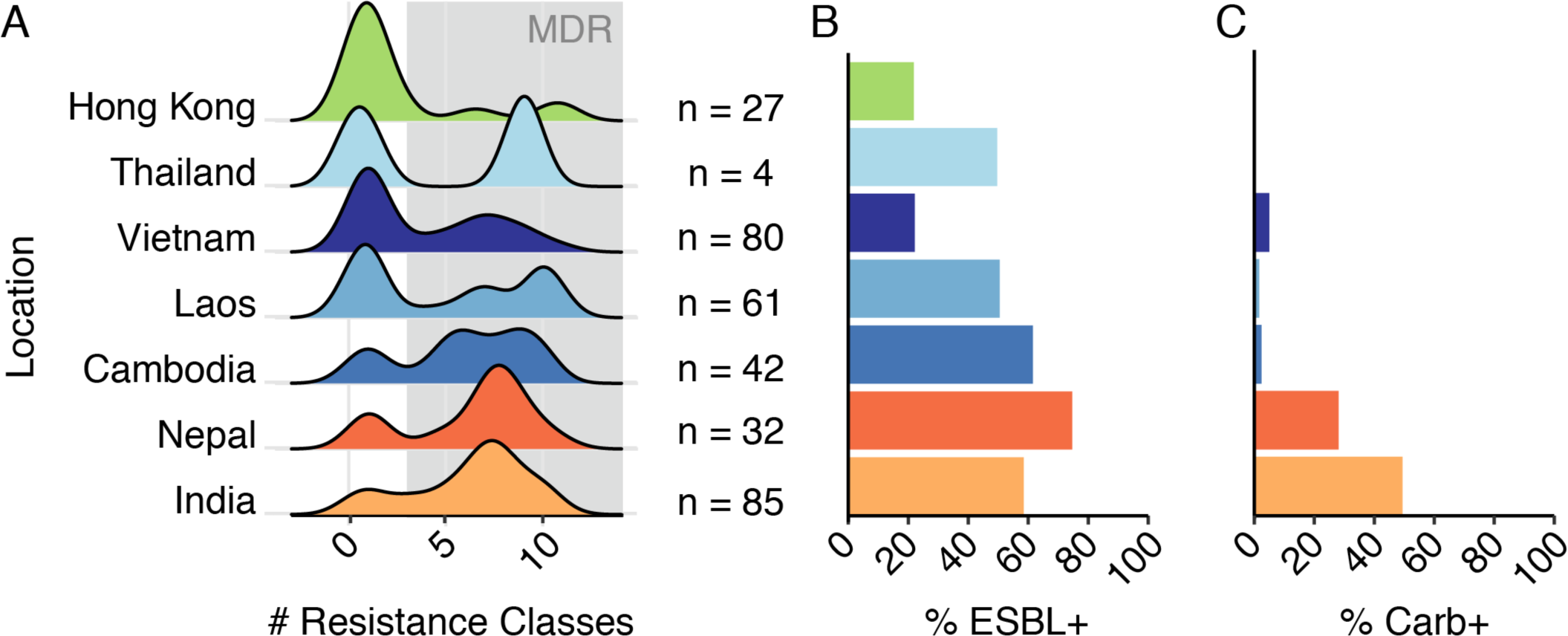
Prevalence of antimicrobial resistance determinants among *K. pneumoniae sensu stricto* isolates. **A)** Density plots showing the distributions of number of drug classes for which acquired resistance determinants were detected in each genome (n = total genomes by location). Grey shading indicates multi-drug resistance (≥3 resistance classes). **B)** Proportion of genomes for which extended-spectrum beta-lactamase genes were detected (% ESBL+). **C)** Proportion of genomes for which carbapenemase genes were detected (% Carb+).

The overall prevalence of ESBL genes amongst the *K. pneumoniae* isolates was 47% (n=157), but varied between study sites (22-75%; see **Figure 3B**). The majority of ESBL *K. pneumoniae* (90%) were predicted to be resistant to a further ≥3 alternative antimicrobial classes (median 7 additional classes). The most common ESBL genes were *bla*_CTX-M-15_ (n=120/157, 76%), *bla*_CTX-M-27_ (n=14/157, 9%) and *bla*_CTX-M-14_ (n=13/157, 8%); these were detected in diverse chromosomal STs (**Table 2, Table S1**). Twelve additional putative ESBL genes were also detected in 1-7 genomes each (see **Table S1**).

**Table 2:** Notable AMR STs by location. Details of all clones with ≥3 ESBL and/or carbapenemase gene positive genomes at any single site. *ST15 is the only clone meeting these criteria at >1 site.

| Location | Total ESBL (%) | Total CP (%) | Notable clone/s | N<sup>a</sup> | N ESBL (% of ESBL/site) | ESBL genes (N) | N CP (% of CP/site) | CP genes (N) |
| --- | --- | --- | --- | --- | --- | --- | --- | --- |
| Hong Kong | 5 (19%) | 0 | - | - | - | - | - | - |
| Vietnam | 20 (25%) | 4 (5%) | * ST15 | 7 | 4 (20%) | CTX-M-15 (3)<br>CTX-M-14 (2) | 2 (50%) | NDM-1 (1) |
| Thailand | 2 (50%) | 0 | - | - | - | - | - | - |
| Laos | 32 (53%) | 1 (2%) | * ST15 | 14 | 14 (44%) | CTX-M-15 (14) | 0 | - |
|  |  |  | ST76-1LV | 4 | 4 (13%) | CTX-M-15 (4) | 0 | - |
| Cambodia | 27 (64%) | 1 (2%) | ST20 | 5 | 5 (19%) | CTX-M-27 (3)<br>CTX-M-14 (2)<br>CTX-M-15 (1) | 0 | - |
|  |  |  | * ST15 | 3 | 3 (11%) | CTX-M-15 (3)<br>CTX-M-27 (1) | 0 | - |
|  |  |  | ST36 | 3 | 3 (11%) | CTX-M-15 (2) | 0 | - |
| Nepal | 24 (75%) | 9 (28%) | * ST15 | 8 | 8 (33%) | CTX-M-15 (8) | 6 (67%) | NDM-1 (5) |
|  |  |  | ST4 | 5 | 5 (21%) | CTX-M-15 (5) | 0 | - |
|  |  |  | ST11 | 3 | 3 (13%) | SFO-1 (3) | 3 (33%) | OXA-232 (3) |
| India | 52 (61%) | 42 (49%) | ST231 | 17 | 13 (25%) | CTX-M-15 (13) | 13 (31%) | OXA-232 (13) |
|  |  |  | ST14 | 16 | 7 (13%) | CTX-M-15 (7) | 11 (26%) | OXA-232 (10) |
|  |  |  | * ST15 | 4 | 3 (6%) | CTX-M-15 (3) | 0 | - |
|  |  |  | ST16 | 3 | 3 (6%) | CTX-M-15 (3) | 2 (5%) | NDM-1 (1) |
|  |  |  | ST29 | 3 | 3 (6%) | CTX-M-15 (3) | 0 | - |
|  |  |  | ST395 | 3 | 1 (25%) | CTX-M-15 (1) | 3 (7%) | OXA-232 (3) |

The overall prevalence of carbapenemase genes was 17% (n=57), again varying widely between sites (0-50%; **Figure 3C**). All isolates with a carbapenemase gene were also MDR, with predicted resistance to a median of six additional drug classes. The most common carbapenemases were the OXA-48-like *bla*_OXA-232_ (n=36/57; 63%) and the metallo- betalactamase *bla*_NDM-1_ (n=18/57; 32%, including four genomes that also carried *bla*_OXA-232_); again these were each detected in a diverse set of STs (**Table 2, Table S1**). Five other carbapenemase genes were detected in 1-4 genomes each (see **Table S1**); *bla*_KPC_ was not detected. Details of ESBL/CP *K. pneumoniae* STs identified at each site, including their specific enzymes, are shown in **Table 2**. Notably ST15 carrying *bla*_CTX-M-15_ were identified at all sites except Thailand (n=4 genomes), and occasionally also harboured the carbapenemases *bla*_NDM-1_ or *bla*_OXA-232_.

### Acquired virulence determinants

Similarly we identified genes linked to invasive disease: the yersiniabactin locus (*ybt*) was present in 163 (49%) of the BSI *K. pneumoniae*, with the site prevalence ranging from 19- 67% (**Figure 4A**); no significant difference in *ybt* prevalence was observed between S and SE Asia (**Table 1**). Nine of the 14 known chromosomally integrated *ybt* mobile-genetic elements (ICE*Kp*s)^15^ and two *ybt* plasmids were detected. The most common were ICE*Kp*5 (n=41), ICE*Kp*4 (n=36) and ICE*Kp*10 (also encoding colibactin, n=31), found across multiple study sites. The ICE*Kp*10-positive samples included 25 in clonal group 23 (ST23 plus related STs), wherein ICE*Kp*10 is a marker of the important globally distributed CG23-I sub-lineage^30^.

**Figure 4:**
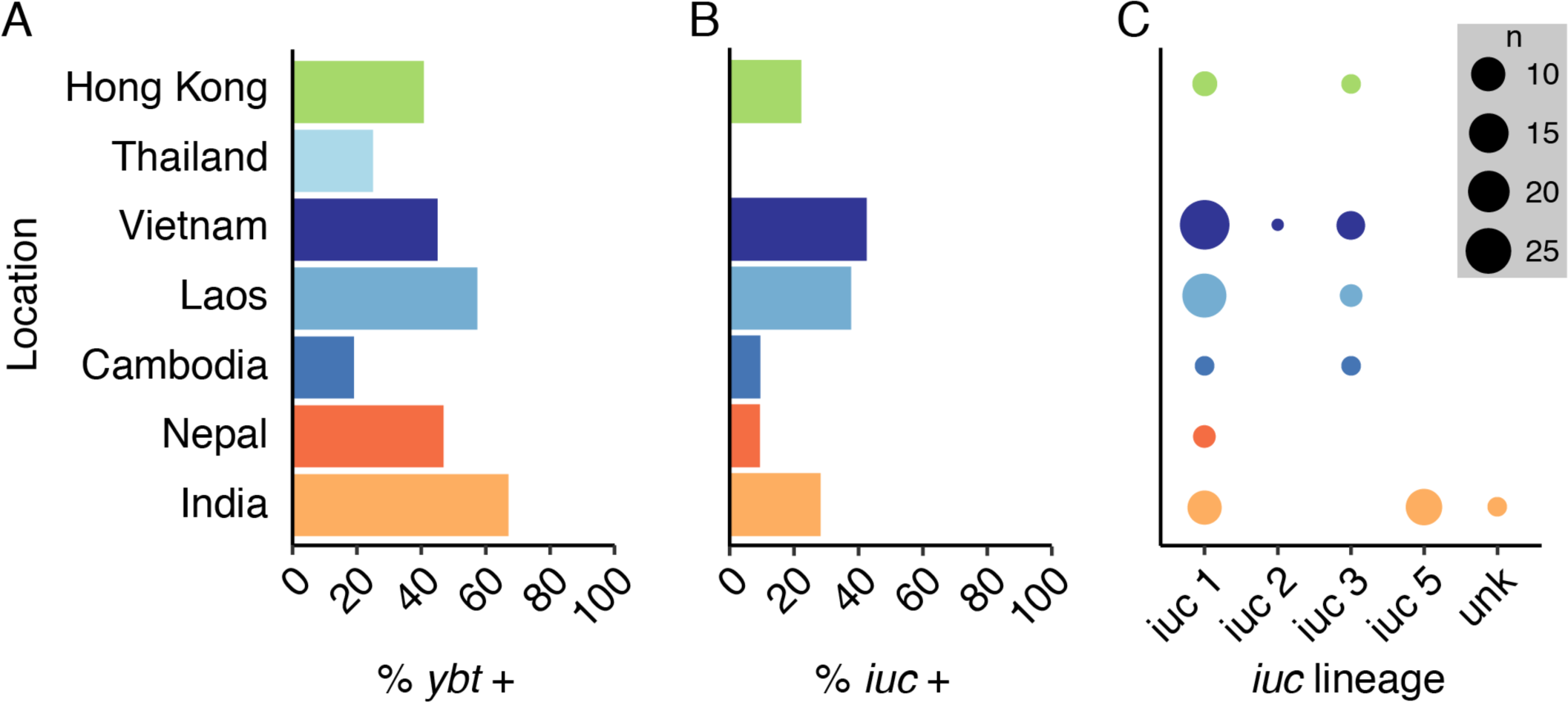
Prevalence of key virulence determinants among *K. pneumoniae sensu stricto* isolates. **A)** Proportion of genomes for which the yersiniabactin locus was detected (% *ybt* +) by location. **B)** Proportion of genomes for which the aerobactin locus was detected (% *iuc* +) by location. **C)** *iuc* lineages by location. Points are scaled by the number of genomes as per legend. Unk; unknown.

The *iuc*, *iro* and *rmpA/rmpA2* loci, typically carried on virulence plasmids, were commonly detected (28% *iuc*, 21% *iro*, 18% *rmpA* 16% *rmpA2*; 14% with all four). The prevalence of *iuc* did not differ significantly between the S and SE Asian isolates, but *iuc* lineages and chromosomal STs of *iuc*-positive isolates were differentially distributed between the sampling sites (**Figure 4B-C, Table 1 and 3**). *Iuc* lineage 1 (*iuc1*, n=66) was widely distributed (**Figure 4C**) but more prevalent in SE Asia (27% vs 13% in S Asia, p=0.011; **Table 1**). *Iuc1* is associated with the KpVP-1 virulence plasmid^31^ and was detected amongst 15 *K. pneumoniae* STs, most commonly those known to be associated with hypervirulent infections: ST23 (n=28), ST65 (n=7) and ST86 (n=7). *Iuc5* (n=12) is associated with *E. coli* plasmids and was detected only in ST231 from India, while *iuc2* (associated with KpVP-2^31^) was detected in a single ST380 isolate from Vietnam. *Iuc3* (n=13) was detected in SE Asia (Cambodia, Vietnam, Laos) and Hong Kong (**Figure 4C**), among eight distinct STs (**Table S1**). We selected four *iuc3*-positive isolates from Vietnam and Laos for long-read sequencing and found each harboured a distinct and novel FIB_K_/FII *iuc3* plasmid (**Table 4**).

### Genotypic convergence of antimicrobial resistance and virulence

While strains carrying either AMR or hypervirulence determinants are of concern, those carrying both pose the greatest potential public health threat. The *ybt* virulence locus was significantly associated with ESBL *K. pneumoniae* (OR 1.7, *p*=0.021) and CP *K. pneumoniae* (OR 3.2, *p*=0.0002). *Ybt*+ ESBL *K. pneumoniae* BSI were detected at all sites (**Figure 5A**) and exhibited a similar prevalence in both S and SE Asia (**Table 1**). *Iuc*, the most significant contributor to hypervirulence in *K. pneumoniae*^17, 32^, was present in 13% of ESBL *K. pneumoniae* BSI (vs 43% amongst non-ESBL; OR 0.2, *p*<0.0001) and 23% of CP *K. pneumoniae* (vs 30% amongst non-CP; OR 0.7, *p*=0.34). Overall prevalence of *iuc*+ ESBL *K. pneumoniae* was 6%. *Iuc*+ ESBL isolates were detected at five of the seven sampling locations (**Figure 5A**), and were more common in S than SE Asia (p<0.001, **Table 1**). Conversely, *iuc*+ CP *K. pneumoniae* BSI were detected only in India (n=12 ST231, 14% of all isolates from this location) and Nepal (n=1 ST15, 3%).

**Figure 5:**
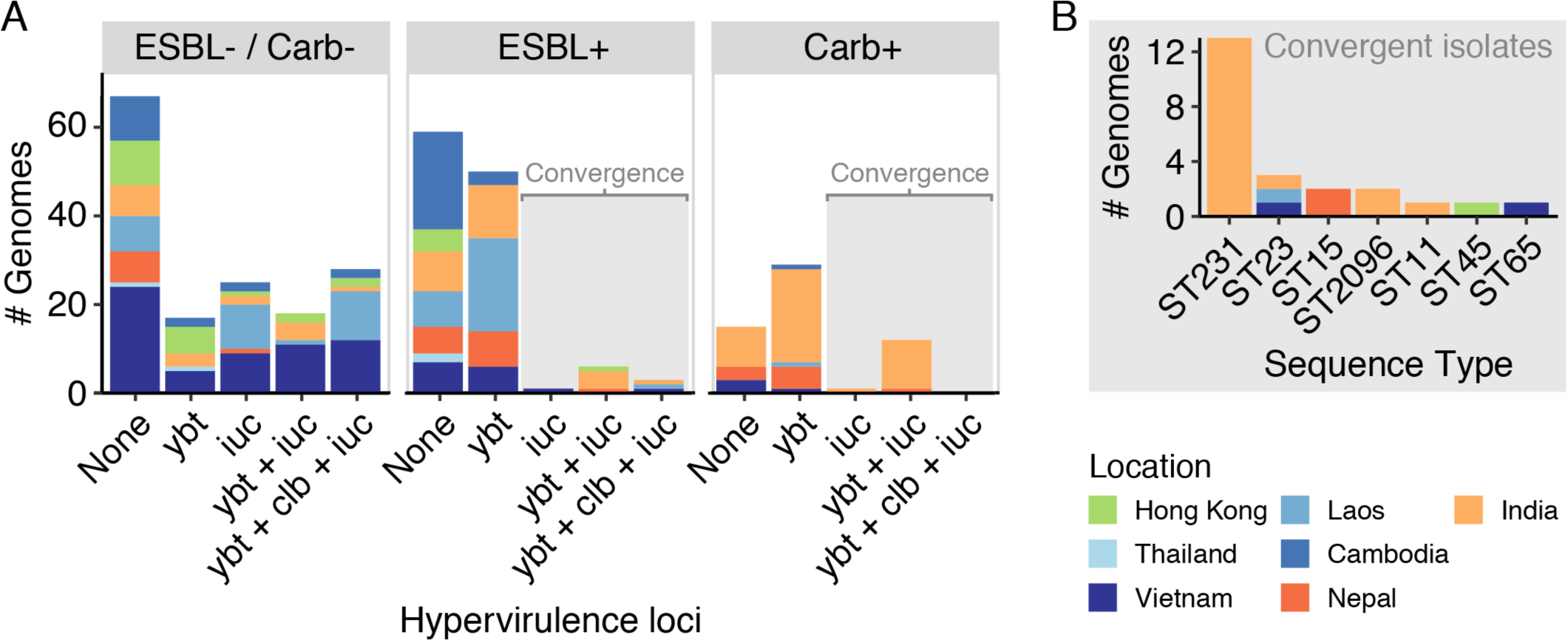
Convergence of virulence and antimicrobial resistance determinants. **A)** Frequency of genomes carrying the yersiniabactin (*ybt*), colibactin (*clb*) and/or aerobactin (*iuc*) loci shown by ESBL and carbapenemase gene status. Bars are coloured by location as per legend. Grey shading indicates convergent isolates i.e. those harbouring at least one ESBL and/or carbapenamse gene plus *iuc* with/without *ybt* and *clb*. **B)** Chromosomal sequence types of convergent isolates. Bars are coloured by location as per legend.

Convergent AMR-hypervirulent isolates (defined as organisms carrying *iuc* in addition to ESBL and/or carbapenemase genes) were seen across seven different STs circulating across this region, with the overall prevalence being 7.3% (**Figure 5B, Table 3**). Long-read sequencing of four representative isolates of different STs revealed four novel mosaic plasmids. Three of these plasmids (and the four *iuc3* plasmids described above) harboured *iuc* plus AMR genes (1-10 AMR genes, encoding resistance to 10 distinct drug classes; see **Table 4**). Notably, one of these plasmids harboured *iuc1*, *bla*_CTX-M-15_, and eight additional AMR genes (plasmid pBA813_1 from isolate BA813, **Table 4).** While pBA813_1 was not predicted to encode the *tra* plasmid transfer machinery, the four *iuc3* plasmids and one of the *iuc1* plasmids (pBA6201_1, *iuc1* plus five AMR genes) contained a complete *tra* operon suggesting that they are capable of conjugative transfer. Consistent with this, we detected a high degree of sequence similarity between the *iuc3* plasmids carried by isolates of three distinct STs in Laos, supporting their dissemination within the local *K. pneumoniae* population.

**Table 3:** Notable iuc-positive STs by region. Details of all STs with ≥3 *iuc*-positive genomes at any single site (i.e. predicted as aerobactin-producing), or carrying *iuc* in addition to ESBL and/or carbapenemase genes. *Known hypervirulent STs. ^a^ICE*Kp*10 carries *ybt* and *clb*.

| Location | Total *iuc*+ | Notable clone/s | *iuc* allele | N *iuc* (% of *iuc*/site) | ICEKp (N) | *iro* allele (N) | *rmpA*/2 (N) | ESBL/CP (N) |
| --- | --- | --- | --- | --- | --- | --- | --- | --- |
| Hong Kong | 6 (22%) | ST45 | *iuc3* | 1 (17%) | ICEKp4 (1) | - | - | CTX-M-3 (1) |
| Vietnam | 34 (43%) | * ST23 | *iuc1* | 13 (38%) | ICEKp10<sup>a</sup> (10)<br>ICEKp3 (1) | *iro1* (13) | (10) | CTX-M-14 (1) |
|  |  | * ST25 | *iuc3* | 2 (6%) | ICEKp1 (2) | *iro3* (2) | (2) | - |
|  |  |  | *iuc1* | 1 (3%) | - | *iro1* (1) | (1) | - |
|  |  | * ST65 | *iuc1* | 2 (6%) | ICEKp10<sup>a</sup> (1) | *iro1* (2) | (2) | CTX-M-15,<br>VEB-1 (1) |
| Thailand | 0 | - | - | - | - | - | - | - |
| Laos | 23 (38%) | * ST23 | *iuc1* | 10 (44%) | ICEKp10<sup>a</sup> (10) | *iro1* (10) | (10) | CTX-M-63 (1) |
|  |  | * ST65 | *iuc1* | 2 (6%) | ICEKp10<sup>a</sup> (2) | *iro1* (3) | (3) | - |
|  |  | * ST86 | *iuc1* | - | ICEKp4 (1) | *iro1* (3) | (3) | - |
|  |  | ST592 | *iuc1* | 10 (44%) | - | *iro1* (3) | (3) | - |
| Cambodia | 4 (10%) | - | - | - | - | - | - | - |
| Nepal | 3 (9%) | ST15 | *iuc1* | 2 (67%) | ICEKp12 (2) | - | (2) | CTX-M-15 (2)<br>OXA-232 (1) |
| India | 24 (28%) | ST231 | *iuc5* | 12 (50%) | ICEKp5 (13) | - | - | CTX-M-15 (10) |
|  |  |  | *iuc*<sup>^</sup> | 2 (8%) |  |  |  |  |
|  |  | ST2096 | *iuc1* | 3 (13%) | ICEKp5 (2) | *iro1* (1) | - | CTX-M-15 (2) |
|  |  | * ST23 | *iuc1* | 2 (8%) | ICEKp3 (1)<br>ICEKp10<sup>a</sup> (1) | *iro1* (2) | (1) | CTX-M-15 (1) |
|  |  | ST11 | *iuc1* | 1 (48%) | ICEKp12 (1) | - | - | CTX-M-15,<br>OXA-232 (1) |

**Table 4:**
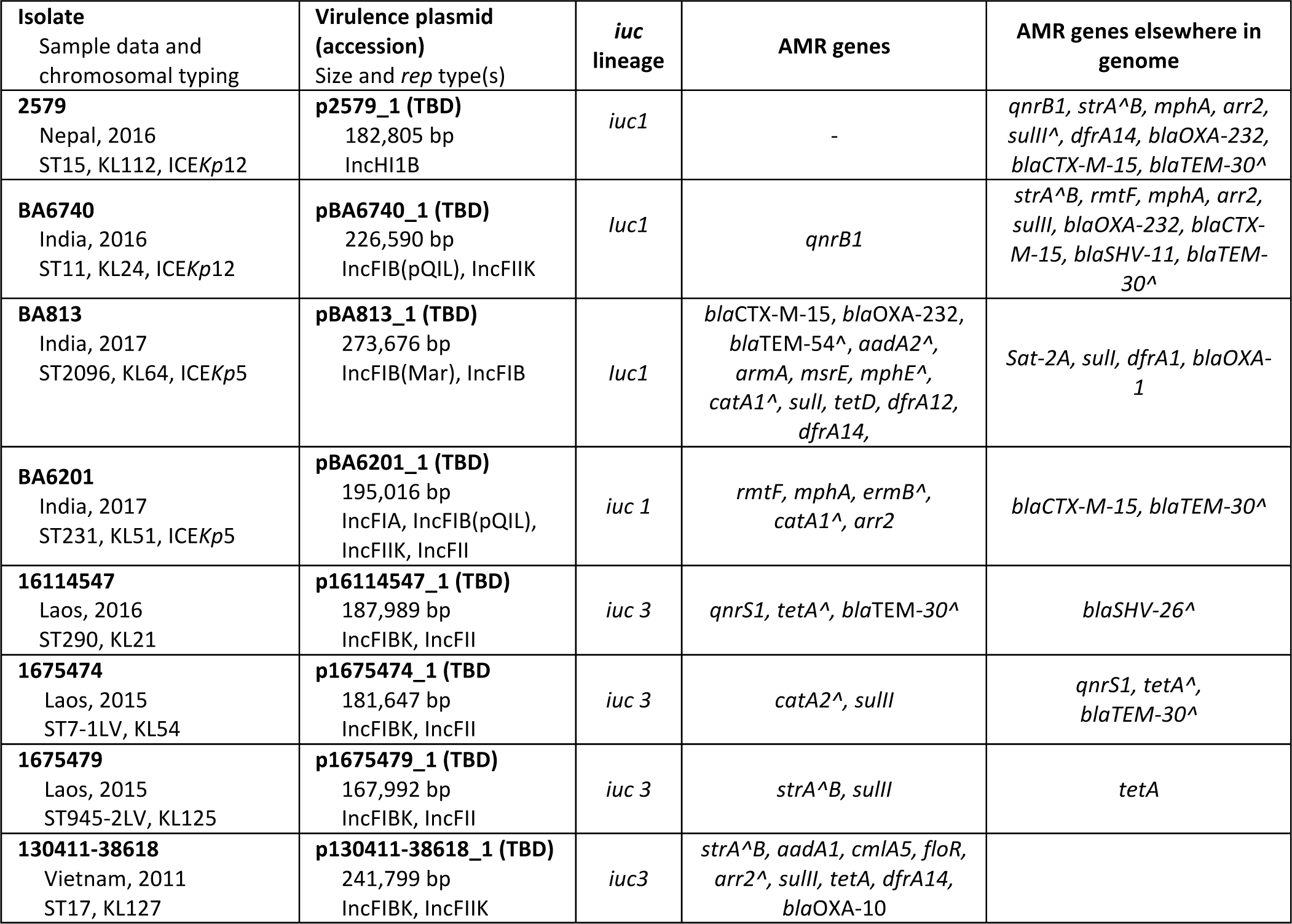
Novel virulence plasmids sequenced in this study and their host isolate properties. *rep* types as per the PlasmidFinder v2.0 database are shown. All plasmids carried the *iuc* aerobactin synthesis operon. No plasmids carried the *iro* or *rmpA/rmpA2* virulence loci. Seven virulence plasmids also carried AMR (antimicrobial resistance) genes. Seven host isolates harboured additional AMR genes elsewhere in the genome. ST; chromosomal multi- locus sequence type. KL; K-locus. ICE*Kp*; integrative conjugative element carrying the yersiniabactin siderophore locus (*ybt*). ^; inexact match.

Comparison of chromosomal ST, *iuc* lineage and AMR gene content indicate at least 9 distinct AMR-virulence convergence events in our sample (**Table 3**). These include four distinct AMR element acquisitions by previously described hypervirulent STs (three in ST23 from Vietnam, Laos and India; one in ST65 from Vietnam), four distinct virulence plasmid acquisitions by previously described MDR STs (one in ST15 from Nepal; one in ST11 from India; one in ST2096 from India; one in ST231 from India; plus one in ST45 from Hong Kong). Most of these organisms also harboured the additional virulence factors *ybt*, *iro* and *rmpA/rmpA2* (**Table 3**), increasing the likelihood of hypervirulent phenotype.

Each AMR-hypervirulent strain was detected only at one site, with no evidence of dissemination between countries. However we identified six clusters of closely related *K. pneumoniae* strains (each separated by ≤100 SNVs) that were detected in different countries (four in multiple SE Asian countries; two in India plus Laos or Hong Kong), suggesting that regional spread of *K. pneumoniae* does occur (**Table S1).**

## Discussion

This work represents the first broad genomic study of *K. pneumoniae* causing BSI in S/SE Asia and Hong Kong, regions that are facing a combination of community-acquired invasive hypervirulent *K. pneumoniae,* unregulated use of antimicrobials, and the emergence and spread of MDR pathogens.

Ominously, alongside the high prevalence of ESBL and carbapenemase genes, our data revealed high prevalence of known hypervirulence determinants with the *iuc* locus detected in 28% of all isolates included in this study, more than double the prevalence seen in previous studies focused outside of this region^31^. While the locus itself was common in both S and SE Asia, the distribution of *iuc* lineages differed (**Table 1**). Specifically, *iuc1* (associated with the characteristic KpVp-1 virulence plasmid^31^) and *iuc3* were more common in SE Asia where they were each detected among numerous distinct STs. This is consistent with their local dissemination, a finding supported by the discovery of four novel *iuc3* encoding plasmids in these isolates. Of greatest concern was the detection of at least 9 distinct convergence events between AMR and hypervirulence (encompassing 7.3% of isolates) including seven novel dual AMR+*iuc* containing plasmids, which is almost twice the total number of convergent plasmids reported so far in other studies^33–36^. Given that our isolate collection represented a snapshot from seven diagnostic laboratories, we predict that these organisms are likely to be more widely distributed across Asia. As such, there is a need for enhanced *K. pneumoniae* surveillance to rapidly identify convergent AMR-hypervirulent strains and/or plasmids, and to monitor their spread in S/SE Asia and beyond.

In the light of these convergence events it is also clear that the diversity of *K. pneumoniae* causing BSI in S/SE Asia represents a significant challenge to therapy using existing or novel agents. Our study revealed a highly diverse set of isolates, both in terms of chromosomal STs and surface polysaccharide loci. The former is consistent with the hypothesis that the majority of BSIs originate from the patients’ personal gastrointestinal microbiota rather than from intra-hospital transmissions^28^. The latter is highly relevant to the design of vaccines or other interventions targeting *K. pneumoniae* capsules or lipopolysaccharide, which are considered of high importance in response to increasing prevalence of ESBL and CP strains, both of which were common in our sampling locations (47% and 17% of all isolates, respectively). From our sampling strategy we crudely estimate a non-cross-reactive capsule- targeted vaccine would need to include ≥16 serotypes in order to provide potential protection against >50% of the BSI *K. pneumoniae* isolates across these study sites. The 16 most common K-loci would cover 61% of all ESBL and 68% of all CP *K. pneumoniae* across S/SE Asia. Alternatively, we estimate that a completely immunising vaccine targeting O1, O2 and O3b would hypothetically protect against 79% of the BSI *K. pneumoniae* isolates in this study; equating to 79% of all ESBL-containing isolates and 70% of CP.

An important limitation of this work is that the data represents a convenience sample of *K. pneumoniae* BSI isolates identified by the participating diagnostic laboratories during routine activities during overlapping time periods, and combined retrospectively. This prevented statistical comparisons between individual sites, and should be considered with caution when extrapolating gene prevalence information to the broader *K. pneumoniae* population. We focussed on BSI, as this allowed for convenient and consistent retrospective identification of isolates associated with invasive disease for inclusion in the study. While this presentation arguably reflects the greatest clinical need, the siderophore virulence loci are known to be more prevalent in BSI compared to other types of infection^14^. In addition, we note that our inferences about AMR are based on genotypic information, which is highly predictive of AMR in *K. pneumoniae* but not perfectly correlated^28, 37^. Nevertheless, our analyses reveal valuable insights, and provide essential data to motivate future systematic studies and enhanced public health surveillance.

Our study represents a blueprint for genomic surveillance of emerging AMR pathogens in Asia and we urge the coordination of similar activities internationally. By rapidly detecting resistance and virulence genes in the context of clonal and surface antigen diversity, our approach provides critical information that can be used to simultaneously track the emergence and dissemination of clinically important variants, guide empirical antimicrobial therapy, and assess potential mechanisms and targets for containment and intervention strategies. In combination with rapid reporting and data sharing, this approach will permit researchers and public health professionals to recognize and control the growing public health threat of AMR *K. pneumoniae*.

## Author contributions

KLW, NRT, SB and KEH conceived the study. DABD, MI, AK, CLL, TM, PNN, LN, AS, PT, BC, MV collected, identified and collated *K. pneumoniae* isolates. TNTN and LMJ generated genome sequence data. KLW, MMCL and KEH analysed the data. KLW and KEH wrote the manuscript. All authors read and commented on the manuscript.

## Acknowledgements

We would like to acknowledge and thank all staff involved in obtaining and processing the isolates including healthcare facility and laboratory staff at Patan Hospital, Kathmandu, Nepal; Christian Medical College Hospital, Vellore, India; Angkor Hospital for Children, Siem Reap, Cambodia; Mahosot Hospital, Vientiane, Lao PDR; The Hospital of Tropical Diseases, Ho Chi Minh City, Vietnam; Prince of Wales Hospital, Hong Kong; and Shoklo Malaria Research Unit, Thailand. We also thank the WSI Pathogen Informatics team for help with data management. This work was supported by the Wellcome Trust (grant #206194 to Wellcome Sanger Institute) and the Bill and Melinda Gates Foundation, Seattle (grant OPP1175797 to KEH). KEH is supported by a Senior Medical Research Fellowship from the Viertel Foundation of Australia. DAB and PNN are supported by the Wellcome Trust.

## SUPPLEMENTARY TABLES

**Table S1 (see spreadsheet): Sample information and genotyping results for all BSI *Kp* included in this study**

**Table S2:** Characteristics of non-*Kp* BSI isolate genomes. ^a^Only O types detected in >1 genome are shown.

|  | *K. quasipneumoniae* subsp. *quasipneumoniae* | *K. quasipneumoniae* subsp. *similipneumoniae* | *K. variicola* |
| --- | --- | --- | --- |
| **N** | 5 | 20 | 9 |
| **# STs** | 4:<br>2 Vietnam<br>1 India<br>1 Nepal | 17:<br>7 Cambodia<br>6 Vietnam<br>5 Laos | 9:<br>3 Hong Kong<br>2 Cambodia<br>1 India<br>1 Laos<br>1 Nepal<br>1 Vietnam |
| **Common STs<sup>a</sup>** | ST1118-3LV (40%) | ST1191 (10%)<br>ST1124 (10%)<br>ST334 (10%) | - |
| **# K loci** | 3 | 15 | 9 |
| **Common K loci<sup>a</sup>** | KL107 (60%) | KL60 (15%)<br>KL10 (10%)<br>KL13 (10%)<br>KL103 (10%) | - |
| **# O types** | 2 | 4 | 4 |
| **Common O types<sup>a</sup>** | O3/O3a (80%) | O5 (55%)<br>O3/O3a (30%)<br>O12 (10%) | O3/O3a (44%)<br>O5 (33%) |
| **% ESBL+** | 0 | 65% | 0 |
| **ESBL genes** | - | CTX-M-15 (35%)<br>CTX-M-27 (10%)<br>SHV-2a (10%)<br>CTX-M-14 (5%)<br>CTX-M-9 (5%) | - |
| **% Carb+** | 20% | 0 | 0 |
| **Virulence determinants** | - | ICEKp3 + *iuc1* + *iro1* + *rmpA*<br>(n=1 ST367 from Vietnam) | *ybt* plasmid lineage<br>(n=1 ST209-1LV from Cambodia) |

**Table S3:**
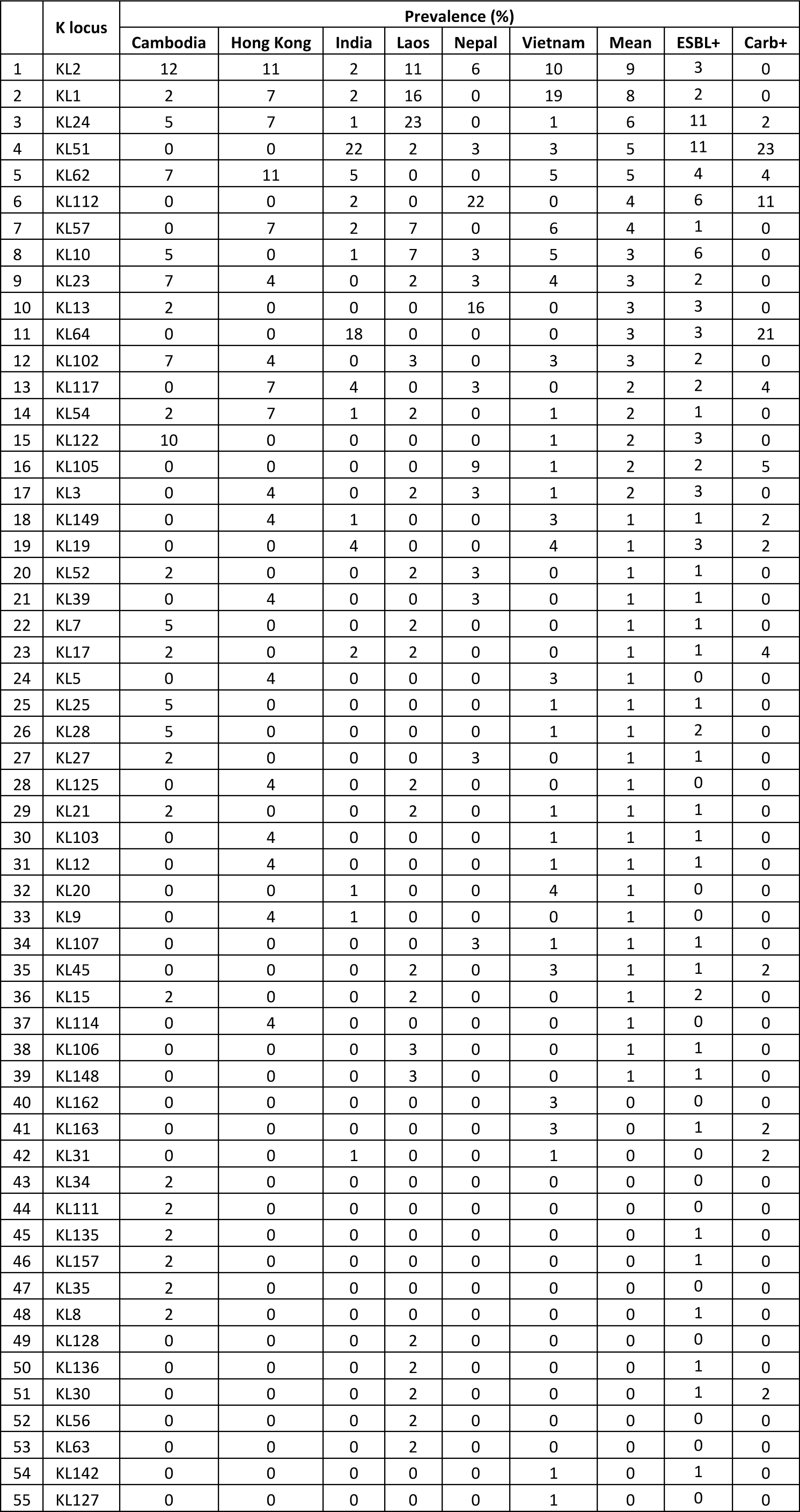
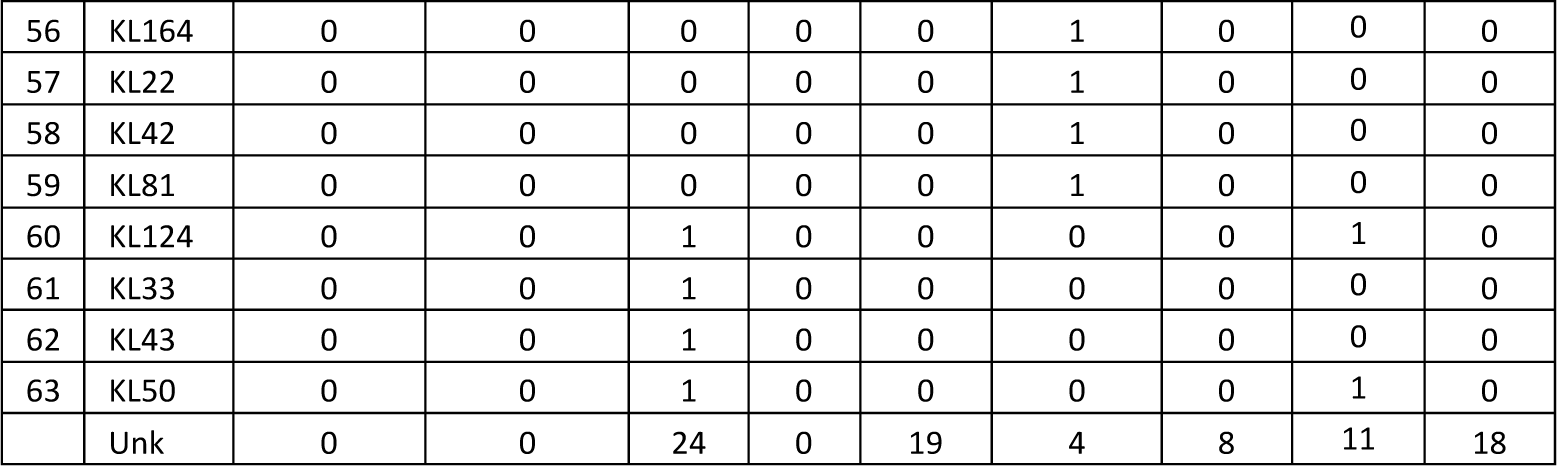
K locus prevalence among K. pneumoniae sensu stricto. K-loci are ordered by highest to lowest mean prevalence across all sites. Note that the Thai site is excluded from these calculations due to small sample size (n=4). Unk; unknown.

|  | K locus | Prevalence (%) |  |  |  |  |  |  |  |  |
| --- | --- | --- | --- | --- | --- | --- | --- | --- | --- | --- |
|  |  | Cambodia | Hong Kong | India | Laos | Nepal | Vietnam | Mean | ESBL+ | Carb+ |
| 1 | KL2 | 12 | 11 | 2 | 11 | 6 | 10 | 9 | 3 | 0 |
| 2 | KL1 | 2 | 7 | 2 | 16 | 0 | 19 | 8 | 2 | 0 |
| 3 | KL24 | 5 | 7 | 1 | 23 | 0 | 1 | 6 | 11 | 2 |
| 4 | KL51 | 0 | 0 | 22 | 2 | 3 | 3 | 5 | 11 | 23 |
| 5 | KL62 | 7 | 11 | 5 | 0 | 0 | 5 | 5 | 4 | 4 |
| 6 | KL112 | 0 | 0 | 2 | 0 | 22 | 0 | 4 | 6 | 11 |
| 7 | KL57 | 0 | 7 | 2 | 7 | 0 | 6 | 4 | 1 | 0 |
| 8 | KL10 | 5 | 0 | 1 | 7 | 3 | 5 | 3 | 6 | 0 |
| 9 | KL23 | 7 | 4 | 0 | 2 | 3 | 4 | 3 | 2 | 0 |
| 10 | KL13 | 2 | 0 | 0 | 0 | 16 | 0 | 3 | 3 | 0 |
| 11 | KL64 | 0 | 0 | 18 | 0 | 0 | 0 | 3 | 3 | 21 |
| 12 | KL102 | 7 | 4 | 0 | 3 | 0 | 3 | 3 | 2 | 0 |
| 13 | KL117 | 0 | 7 | 4 | 0 | 3 | 0 | 2 | 2 | 4 |
| 14 | KL54 | 2 | 7 | 1 | 2 | 0 | 1 | 2 | 1 | 0 |
| 15 | KL122 | 10 | 0 | 0 | 0 | 0 | 1 | 2 | 3 | 0 |
| 16 | KL105 | 0 | 0 | 0 | 0 | 9 | 1 | 2 | 2 | 5 |
| 17 | KL3 | 0 | 4 | 0 | 2 | 3 | 1 | 2 | 3 | 0 |
| 18 | KL149 | 0 | 4 | 1 | 0 | 0 | 3 | 1 | 1 | 2 |
| 19 | KL19 | 0 | 0 | 4 | 0 | 0 | 4 | 1 | 3 | 2 |
| 20 | KL52 | 2 | 0 | 0 | 2 | 3 | 0 | 1 | 1 | 0 |
| 21 | KL39 | 0 | 4 | 0 | 0 | 3 | 0 | 1 | 1 | 0 |
| 22 | KL7 | 5 | 0 | 0 | 2 | 0 | 0 | 1 | 1 | 0 |
| 23 | KL17 | 2 | 0 | 2 | 2 | 0 | 0 | 1 | 1 | 4 |
| 24 | KL5 | 0 | 4 | 0 | 0 | 0 | 3 | 1 | 0 | 0 |
| 25 | KL25 | 5 | 0 | 0 | 0 | 0 | 1 | 1 | 1 | 0 |
| 26 | KL28 | 5 | 0 | 0 | 0 | 0 | 1 | 1 | 2 | 0 |
| 27 | KL27 | 2 | 0 | 0 | 0 | 3 | 0 | 1 | 1 | 0 |
| 28 | KL125 | 0 | 4 | 0 | 2 | 0 | 0 | 1 | 0 | 0 |
| 29 | KL21 | 2 | 0 | 0 | 2 | 0 | 1 | 1 | 1 | 0 |
| 30 | KL103 | 0 | 4 | 0 | 0 | 0 | 1 | 1 | 1 | 0 |
| 31 | KL12 | 0 | 4 | 0 | 0 | 0 | 1 | 1 | 1 | 0 |
| 32 | KL20 | 0 | 0 | 1 | 0 | 0 | 4 | 1 | 0 | 0 |
| 33 | KL9 | 0 | 4 | 1 | 0 | 0 | 0 | 1 | 0 | 0 |
| 34 | KL107 | 0 | 0 | 0 | 0 | 3 | 1 | 1 | 1 | 0 |
| 35 | KL45 | 0 | 0 | 0 | 2 | 0 | 3 | 1 | 1 | 2 |
| 36 | KL15 | 2 | 0 | 0 | 2 | 0 | 0 | 1 | 2 | 0 |
| 37 | KL114 | 0 | 4 | 0 | 0 | 0 | 0 | 1 | 0 | 0 |
| 38 | KL106 | 0 | 0 | 0 | 3 | 0 | 0 | 1 | 1 | 0 |
| 39 | KL148 | 0 | 0 | 0 | 3 | 0 | 0 | 1 | 1 | 0 |
| 40 | KL162 | 0 | 0 | 0 | 0 | 0 | 3 | 0 | 0 | 0 |
| 41 | KL163 | 0 | 0 | 0 | 0 | 0 | 3 | 0 | 1 | 2 |
| 42 | KL31 | 0 | 0 | 1 | 0 | 0 | 1 | 0 | 0 | 2 |
| 43 | KL34 | 2 | 0 | 0 | 0 | 0 | 0 | 0 | 0 | 0 |
| 44 | KL111 | 2 | 0 | 0 | 0 | 0 | 0 | 0 | 0 | 0 |
| 45 | KL135 | 2 | 0 | 0 | 0 | 0 | 0 | 0 | 1 | 0 |
| 46 | KL157 | 2 | 0 | 0 | 0 | 0 | 0 | 0 | 1 | 0 |
| 47 | KL35 | 2 | 0 | 0 | 0 | 0 | 0 | 0 | 0 | 0 |
| 48 | KL8 | 2 | 0 | 0 | 0 | 0 | 0 | 0 | 1 | 0 |
| 49 | KL128 | 0 | 0 | 0 | 2 | 0 | 0 | 0 | 0 | 0 |
| 50 | KL136 | 0 | 0 | 0 | 2 | 0 | 0 | 0 | 1 | 0 |
| 51 | KL30 | 0 | 0 | 0 | 2 | 0 | 0 | 0 | 1 | 2 |
| 52 | KL56 | 0 | 0 | 0 | 2 | 0 | 0 | 0 | 0 | 0 |
| 53 | KL63 | 0 | 0 | 0 | 2 | 0 | 0 | 0 | 0 | 0 |
| 54 | KL142 | 0 | 0 | 0 | 0 | 0 | 1 | 0 | 1 | 0 |
| 55 | KL127 | 0 | 0 | 0 | 0 | 0 | 1 | 0 | 0 | 0 |
| 56 | KL164 | 0 | 0 | 0 | 0 | 0 | 1 | 0 | 0 | 0 |
| 57 | KL22 | 0 | 0 | 0 | 0 | 0 | 1 | 0 | 0 | 0 |
| 58 | KL42 | 0 | 0 | 0 | 0 | 0 | 1 | 0 | 0 | 0 |
| 59 | KL81 | 0 | 0 | 0 | 0 | 0 | 1 | 0 | 0 | 0 |
| 60 | KL124 | 0 | 0 | 1 | 0 | 0 | 0 | 0 | 1 | 0 |
| 61 | KL33 | 0 | 0 | 1 | 0 | 0 | 0 | 0 | 0 | 0 |
| 62 | KL43 | 0 | 0 | 1 | 0 | 0 | 0 | 0 | 0 | 0 |
| 63 | KL50 | 0 | 0 | 1 | 0 | 0 | 0 | 0 | 1 | 0 |
|  | Unk | 0 | 0 | 24 | 0 | 19 | 4 | 8 | 11 | 18 |

**Table S4:** Predicted O type prevalence among *K. pneumoniae sensu stricto.* Predicted O types are ordered by highest to lowest mean prevalence across all sites. Note that the Thai site is excluded from these calculations due to small sample size (n=4). ^a^ It was not possible to confidently distinguish between O1 and O2 for a minority of genomes (n=12 in total). The prevalence of these genomes is shown here by site and among ESBL+/Carb+ isolates. These values are added at the second position in the cumulative prevalence plot (**Figure 2B**). Unk; unknown.

| O type | Prevalence (%) |  |  |  |  |  |  |  |  |
| --- | --- | --- | --- | --- | --- | --- | --- | --- | --- |
|  | Cambodia | Hong Kong | India | Laos | Nepal | Vietnam | Mean | ESBL+ | Carb+ |
| O1 | 67 | 48 | 35 | 69 | 66 | 50 | 56 | 55 | 44 |
| O2 | 12 | 26 | 15 | 8 | 16 | 15 | 15 | 13 | 16 |
| O1/O2<sup>a</sup> | 0 | 4 | 11 | 0 | 0 | 3 | 3 | 5 | 7 |
| O3b | 10 | 11 | 6 | 10 | 3 | 9 | 8 | 6 | 4 |
| O4 | 2 | 4 | 0 | 2 | 3 | 4 | 2 | 3 | 0 |
| O5 | 7 | 4 | 0 | 0 | 0 | 4 | 2 | 1 | 0 |
| O3/O3a | 0 | 0 | 0 | 8 | 0 | 3 | 2 | 3 | 0 |
| OL101 | 2 | 0 | 0 | 0 | 6 | 1 | 2 | 1 | 0 |
| OL103 | 0 | 4 | 0 | 2 | 0 | 1 | 1 | 1 | 0 |
| OL104 | 0 | 0 | 2 | 0 | 0 | 0 | 0 | 1 | 2 |
| OL102 | 0 | 0 | 0 | 0 | 0 | 1 | 0 | 0 | 0 |
| Unk | 0 | 0 | 31 | 2 | 6 | 10 | 8 | 12 | 28 |

## References

1. World Health Organization. Global priority list of antibiotic-resistant bacteria to guide research, discovery, and devlopment of new antibiotics. 2017.

2. Rello J, Kalwaje Eshwara V, Lagunes L, et al. A global priority list of the TOp TEn resistant Microorganisms (TOTEM) study at intensive care: a prioritization exercise based on multi-criteria decision analysis. Eur J Clin Microbiol Infect Dis 2018; Advanced online publication.

3. Dat VQ, Vu HN, Nguyen The H, et al. Bacterial bloodstream infections in a tertiary infectious diseases hospital in Northern Vietnam: aetiology, drug resistance, and treatment outcome. BMC Infect Dis 2017; 17: 493.

4. Fox-Lewis A, Takata J, Miliya T, et al. Antimicrobial resistance in invasive bacterial infections in hospitalized children, Cambodia, 2007–2016. Emerg Infect Dis 2018; 24: 841–51.

5. Anderson M, Luangxay K, Sisouk K, et al. Epidemiology of bacteremia in young hospitalized infants in Vientiane, Laos, 2000-2011. J Trop Pediatr 2014; 60: 10–6.

6. The HC, Karkey A, Thanh DP, et al. A high-resolution genomic analysis of multidrug- resistant hospital outbreaks of *Klebsiella pneumoniae*. EMBO Molec Med 2015; 7: 227–39.

7. Hsu LY, Tan TY, Jureen R, et al. Antimicrobial drug resistance in Singapore hospitals. Emerg Infect Dis 2007; 13: 1944–7.

8. Hamzan NI, Yean CY, Rahman RA, Hasan H, Rahman ZA. Detection of *bla*IMP4 and *bla*NDM1 harboring *Klebsiella pneumoniae* isolates in a university hospital in Malaysia. Emerg Health Threats J 2015; 8: 8–12.

9. Jajoo M, Manchanda V, Chaurasia S, et al. Alarming rates of antimicrobial resistance and fungal sepsis in outborn neonates in North India. PLoS One 2018; 13: e0180705.

10. Mohanty S, Gajanand M, Gaind R. Identification of carbapenemase-mediated resistance among *Enterobacteriaceae* bloodstream isolates: A molecular study from India. Indian J Med Microbiol 2017; 35: 421.

11. Smit P, Stoesser N, Pol S, et al. Transmission dynamics of hyper-endemic multi-drug resistant *Klebsiella pneumoniae* in a Southeast Asian neonatal unit: a longitudinal study with whole genome sequencing. Front Microbiol 2018; 9: 1197.

12. Castanheira M, Deshpande LM, Mathai D, Bell JM, Jones RN, Mendes RE. Early dissemination of NDM-1- and OXA-181-producing Enterobacteriaceae in Indian hospitals: Report from the SENTRY Antimicrobial Surveillance Program, 2006-2007. Antimicrob Agents Chemother 2011; 55: 1274–8.

13. Martin RM, Bachman MA. Colonization, infection, and the accessory genome of *Klebsiella pneumoniae*. Front Cell Infect Microbiol 2018; 8: 1–15.

14. Holt KE, Wertheim H, Zadoks RN, et al. Genomic analysis of diversity, population structure, virulence, and antimicrobial resistance in *Klebsiella pneumoniae*, an urgent threat to public health. Proc Natl Acad Sci U S A 2015; 112: E3574-81.

15. Lam MMC, Wick RR, Wyres KL, et al. Genetic diversity, mobilisation and spread of the yersiniabactin-encoding mobile element ICE*Kp* in *Klebsiella pneumoniae* populations. MGen 2018; 4. DOI:http://dx.doi.org/10.1101/098178.

16. Siu LK, Yeh KM, Lin JC, Fung CP, Chang FY. *Klebsiella pneumoniae* liver abscess: A new invasive syndrome. Lancet Infect Dis 2012; 12: 881–5.

17. Russo TA, Olson R, Fang C-T, et al. Identification of biomarkers for differentiation of hypervirulent *Klebsiella pneumoniae* from classical *K. pneumoniae*. J Clin Microbiol 2018; 56: e00776-18.

18. Gu D, Dong N, Zheng Z, et al. A fatal outbreak of ST11 carbapenem-resistant hypervirulent *Klebsiella pneumoniae* in a Chinese hospital: a molecular epidemiological study. Lancet Infect Dis 2017; 3099: 1–10.

19. Wick RR, Judd LM, Gorrie CL, Holt KE. Completing bacterial genome assemblies with multiplex MinION sequencing. MGen 2017; 3. DOI:10.1101/160614.

20. Bankevich A, Nurk S, Antipov D, et al. SPAdes: a new genome assembly algorithm and its applications to single-cell sequencing. J Comput Biol 2012; 19: 455–77.

21. Wick RR, Judd LM, Gorrie CL, Holt KE. Unicycler: resolving bacterial genome assemblies from short and long sequencing reads. PLoS Comp Biol 2017; 13: e1005595.

22. Wyres KL, Wick RR, Gorrie C, et al. Identification of *Klebsiella* capsule synthesis loci from whole genome data. Microb Genomics 2016; 2. DOI:10.1099/mgen.0.000102.

23. Wick RR, Heinz E, Holt KE, Wyres KL. Kaptive Web: user-friendly capsule and lipopolysaccharide serotype prediction for *Klebsiella* genomes. J Clin Microbiol 2018; 56: e00197-18.

24. Wick RR, Schultz MB, Zobel J, Holt KE. Bandage: interactive visualization of *de novo* genome assemblies. Bioinformatics 2015; 31: 3350–2.

25. Seemann T. Prokka: rapid prokaryotic genome annotation. Bioinformatics 2014; 30: 2068–9.

26. Harris SR. SKA: Split Kmer Analysis Toolkit for bacterial genomic epidemiology. bioRxiv 2018;: bioRxiv preprint.

27. Carattoli A, Zankari E, Garciá-Fernández A, et al. PlasmidFinder and pMLST: *in silico* detection and typing of plasmids. Antimicrob Agents Chemother 2014; 58: 3895–903.

28. Gorrie CL, Mirceta M, Wick RR, et al. Gastrointestinal carriage is a major reservoir of *K. pneumoniae* infection in intensive care patients. Clin Infect Dis 2017; Epub ahead of print.

29. Follador R, Heinz E, Wyres KL, et al. The diversity of *Klebsiella pneumoniae* surface polysaccharides. Microb Genomics 2016; 2. DOI:10.1099/mgen.0.000073.

30. Lam MMC, Wyres KL, Duchêne S, et al. Population genomics of hypervirulent *Klebsiella pneumoniae* clonal group 23 reveals early emergence and rapid global dissemination. Nat Commun 2018; 9: 2703.

31. Lam MCC, Wyres KL, Judd LM, et al. Tracking key virulence loci encoding aerobactin and salmochelin siderophore synthesis in *Klebsiella pneumoniae*. Genome Med 2018; 10: 77.

32. Russo TA, Olson R, MacDonald U, et al. Aerobactin mediates virulence and accounts for increased siderophore production under iron-limiting conditions by hypervirulent (hypermucoviscous) *Klebsiella pneumoniae*. Infect Immun 2014; 82: 2356–67.

33. Turton JF, Payne Z, Coward A, et al. Virulence genes in isolates of Klebsiella pneumoniae from the UK during 2016, including among carbapenemase gene- positive hypervirulent K1-ST23 and ‘non-hypervirulent’ types ST147, ST15 and ST383. J Med Microbiol 2017; 67: 118–28.

34. Dong N, Lin D, Zhang R, Chan EWC, Chen S. Carriage of *bla*KPC-2 by a virulence plasmid in hypervirulent *Klebsiella pneumoniae*. J Antimicrob Chemother 2018; 73: 3317–21.

35. Lam MMC, Wyres KL, Wick RR, et al. Convergence of virulence and multidrug resistance in a single plasmid vector in multidrug-resistant *Klebsiella pneumoniae* ST15. J Antimicrob Chemother 2019. Advanced online publication.

36. Shen D, Ma G, Li C, et al. Emergence of a multidrug-resistant hypervirulent *Klebsiella pneumoniae* of ST23 with a rare *bla_CTX-M-24_*-harboring virulence plasmid. Antimicrob Agents Chemother 2019; Advanced online publication.

37. Stoesser N, Batty EM, Eyre DW, et al. Predicting antimicrobial susceptibilities for *Escherichia coli* and *Klebsiella pneumoniae* isolates using whole genomic sequence data. J Antimicrob Chemother 2013; 68: 2234–44.

